# MicroRNA-221/222-expression in HSC and MPP safeguards their quiescence and multipotency by downregulating stress-independent and dependent expression of IEG and of several myelo/granulopoiesis-enhancing target genes

**DOI:** 10.1101/2023.01.30.526397

**Authors:** Peter K. Jani, Georg Petkau, Yohei Kawano, Uwe Klemm, Gabriela Maria Guerra, Gitta Anne Heinz, Frederik Heinrich, Pawel Durek, Mir-Farzin Mashreghi, Fritz Melchers

**Affiliations:** Deutsches Rheuma Forschungszentrum (DRFZ), Berlin, Germany; Max Planck Institute for Infection Biology, Berlin, Germany

## Abstract

The microRNA cluster-221/222 is expressed in hematopoietic stem cells (HSC) and multipotent progenitors (MPP). To study its function in hematopoiesis, we generated mice, in which this cluster is selectively deleted by Vav-cre in HSC and, thus, in all hematopoietic cells. Fluorescence-activated cell sorting analyses of the lineage-negative HSC and MPP compartments in bone marrow at unperturbed, steady state hematopoiesis detect strong activation of HSC to MPP, as well as increased granulocytes in the periphery, induced by miR-221/222-deficiency. Short-term social stress on mice also activates HSC to MPP, but the time of stress is too short to detect further increases in granulocyte numbers. Single cell deep mRNA sequencing identifies Fos as direct, and Jun as well as six other immediate early genes (IEG) as indirect targets of miR-221/222 at unperturbed hematopoiesis. Three of these IEG - Klf6, Nr4a1 and Zfp36 - have previously been found to influence myelo/granulopoiesis. Short stress induces higher levels of the same, and an even larger number of IEGs, now also in MPP, indicating, that stress and miR-221/222 both activate HSC to MPP by IEG upregulation in perturbed hematopoiesis. Furthermore, combined stress and miR-221/222-deficiency rapidly increase numbers of myelo/granulocyte progenitors (MEP, GMP) in bone marrow. Additional indirect miR-221/222-targets become detectable in MPP, of which H3f3b has previously been found to influence myelopoiesis. In serial transplantations, miR-221/222-deficient HSC retain their capacity to home to, and become resident in bone marrow, but they loose their lymphopoietic capacities, thus their multipotency. Our results suggest, that miR-221/222-expression in HSC and MPP safeguards their quiescence and multipotency by downregulating the expression of IEG and of myelo/granulopoiesis-enhancing target genes. Since miR-221/222 is also expressed in human HSC and MPP, its expression should improve clinical settings of human bone marrow transplantations.

## INTRODUCTION

Throughout life, hematopoietic stem cells (HSC) reside in bone marrow (BM), from where they continue to generate, at steady, unperturbed state, all hematopoietic cell lineages(1–7). When transplanted, HSC reconstitute all hematopoietic stem and progenitor cell compartments and all hematopoietic cell lineages in normal numbers in the host. Upon activation, HSC enter cell cycle, and, when differentiating, give rise to multipotent progenitors (MPP1-MPP4) which generate common lymphoid (CLP) and to common myeloid progenitors (CMP). HSC are heterogeneous, with multilineage-potential, myeloid-erythroid-biased, myeloid-erythroid-restricted, and differentiation-inactive HSC(3, 8, 9). How this heterogeneity is regulated, is poorly understood.

Hematopoiesis can be perturbed by different forms of stress(1, 4, 10–24). Here, we investigate three forms of stress. First, before “ex vivo/in vitro” preparation of BM cells for FACS analyses a 20-hour-long transportation of the mice from the breeding facility to the laboratory induces social stress(12). This social stress is reversible by a 7-day-rest period as it is recommended after housing mice in a new environment. Second, during “ex vivo/in vitro” cell preparation BM cells become exposed to “in vitro” oxidative(10) and shear(11) stress. We inhibit the shorter influence of “in vitro” stress on transcription by the addition of the RNA polymerase inhibitor Actinomycin D (ActD) during “ex vivo/in vitro” BM cell preparations for all transcriptional analyses(25–27). Third, “ex vivo/in vitro” stress is probably only one of several forms of stress, which impact HSC, when they are transplanted repeatedly in serial transplantations to study their homing and reconstitution potential(1, 4). We compare pool sizes of HSC and of progenitor compartments (MPP1 and 2), as well as single cell transcriptomic describing their gene expression programs, unperturbed at steady state with those perturbed by either social stress, “in vitro” stress or by transplantation. In BM, social stress reduces numbers of HSC approximately three-fold and increased numbers of MPPs one and a half-fold. Single cell transcriptome analyses of socially or of “in vitro” stressed HSC and MPPs identified more than 70 stress-dependently upregulated genes in each form of stress, among them at least seven Fos/Jun AP-1 transcription factor-controlled immediate early genes (IEG) in both forms of stress.

MicroRNAs (miRNA) as post-transcriptional regulators of gene expression have been shown to play a crucial role in hematopoiesis, modulating single or multiple genes in multiple cellular functions(28–33). Specific ablation of Dicer in HSCs, thus complete deletion of mature miRNAs, leads to increased apoptosis in HSCs and affects their self-renewal capacity(33–35). Several miRNAs have been identified, which influence the regulations of these different states of HSC(34, 36) by the mTOR pathway(37, 38).

The cluster of miR-221 and miR-222 was found to be expressed in early hematopoietic progenitors(38, 39). Except for a previous report which indicated a potential involvement of the miR-221/222 cluster in early erythropoiesis the role of this microRNA cluster in early hematopoiesis remained unknown(40). Furthermore, we have previously described the influences of miR-221 on preBI-cell homing to BM(41). MiR-221 overexpression confers the ability to these cells to home to, and become resident in special, subosteal areas in the bone marrow after transplantation. This is accomplished by activation of the PI3K signaling network in response to BM niche factors like the chemokine CXCL12. This activation induces integrin VLA-4 to change to its high affinity binding conformation, which allows increased adhesion and residence in environments of BM expressing the cell adhesion molecule VCAM1(41). Since HSCs have the natural ability to home to and engraft in BM, we reasoned, that miR221/222 might influence HSC pools in BM by regulating their engraftment as in preB cells. As we show here, this hypothesis is not tenable for HSC.

Here, we first measure the numbers of miR-221 and miR-222 molecules expressed in single HSC, MPP1 and MPP2, and show, that all HSC and MPP express the miRNA cluster. We then generate mice, in which HSC and their subsequent MPPs and hematopoietic lineages become targetable for deletion of the floxed miR-221/222 cluster, using Vav-promotor-induced expression of iCre(9, 42). Single cell expression analyses show that all HSC and MPPs have their miR-221/222 cluster deleted.

Deletion of the miR-221/222 cluster results in reduced numbers of HSC during steady state hematopoiesis, accompanied by increased numbers of multipotent progenitors (MPP1-4) similar to the effect of social stress perturbation in miR221/222-proficien mice. Importantly, the number of granulocytes in spleen is increased by the miR-221/222-deficiency.

In addition, miR-221/222-deficiency upon social stress perturbation selectively increases numbers of granulocyte progenitors in BM. In order to understand the molecular mechanisms of HSC actions and identify direct miR-221/222 targets, we generate a single cell transcriptional landscape of hematopoietic progenitors in BM. The differential single cell transcriptome analyses of unperturbed, miR-221/222-proficient with deficient HSC and MPPs identify Fos as direct miRNA target. Together with increased levels of *Jun*, *Fos* forms increased amounts of the heterodimeric activator protein-1 (AP-1)(43). Furthermore, five other immediate early response genes (IEG), Ier2, Junb, Klf6, Nr4a1 and Zfp36, like Fos and Jun all AP-1-controlled(44), are selectively increased in expression as indirect target genes. The comparisons of single cell mRNA-deep sequencing analyses of socially stressed with “in vitro” stressed, and miR-221/222-deficient HSC identify five of the seven Fos/AP-1-controlled IEG genes, Ier2, Jun, Junb, Klf6 and Zfp36 as common activators of HSC from quiescence. This miR-221/222-deficiency-induced Fos/AP-1 activation became only detectable, because we were able to control the social stress-and “in vitro”-stress-mediated perturbations induced in HSC and progenitors.

Single cell transcriptomes of socially stressed, miR-221/222-deficient HSC and MPPs identified miR-221/222-deficiency-dependent, stressed expression of genes encoding heat shock proteins (Hspa5 and 8), tubulin-cytoskeleton-organizing proteins (Tuba1b, Tubb 4b and 5) and chromatin remodeling proteins (H3f3b, H2afx, H2afz, Hmgb2), upregulated in HSC, MPP1 and/or MPP2, as potential regulators of stress-induced, miR-221/222-dependent increased granulocyte differentiation. Finally, combined stress induced by serial transplantations and miR-221/222-deficiency exhausts lymphoid differentiation, but not BM-homing and granulocyte-differentiation capacities of HSC.

Thus, our findings identify the miR-221/222 cluster as regulator of Fos/AP-1-IEG responses of HSC quiescence and multipotency, but not of HSC engraftment in BM. The results of our studies presented here suggest that expression of miR-221/222 safeguards HSC pools from depletion(9, 45) and maintains their multipotency(21, 24, 46, 47), findings, which have direct implications for improved bone marrow transplant management.

## RESULTS

### Controlled perturbations of steady-state hematopoiesis can be induced by “in vivo” social stress or by “ex vivo/in vitro” oxidative/shear stress

Throughout life, all hematopoietic cell lineages are continuously renewed at steady state from HSC(1–7). Steady-state hematopoiesis can be perturbed by various forms of stress(1, 4, 10–24), leading to increased activation of HSC from quiescence. In order to study the influences of a miRNA cluster, miR-221/222 on the sizes of hematopoietic progenitor compartments in BM and their gene expression activities, we aimed at controlling the states of either non-perturbed, steady-state or perturbed hematopoiesis. Perturbations by social stress(12) or by “ex vivo/in vitro” oxidative/shear stress(10, 11) are experimentally controllable. A 20 hour-long social stress during mouse transportations can be reversed by a subsequent 7 days-rest period, while the initial status of “ex vivo/in vitro” single cell mRNA expressions, analyzed by single cell transcriptome analyses, can be preserved in HSC, MPPs and all other hematopoietic cells of BM by the inhibition of RNA polymerase at the start of BM cell isolation for “in vitro” analyses.

### Controlled perturbation of steady-state hematopoiesis by “in vivo” social stress reduces HSC pools and increases MPPs in BM

Social stress imposed for 20 hours permitted us to measure, and to compare the actions of social stress on pool sizes of miR-221/222-proficient with deficient BM cells. We found, that social stress reduced the HSC pool in BM two-fold and increased MPP2 two-and-a-half-fold. (**Fig 1A**). The numbers of MPP3, MPP4, CLP, CMP and peripheral granulocytes, T and B cells were not affected by short term stress (**Supplementary Fig 1.** For nomenclature see Wilson et al. and Cabezas-Wallscheid et al.(5, 7)).

**Fig 1:**
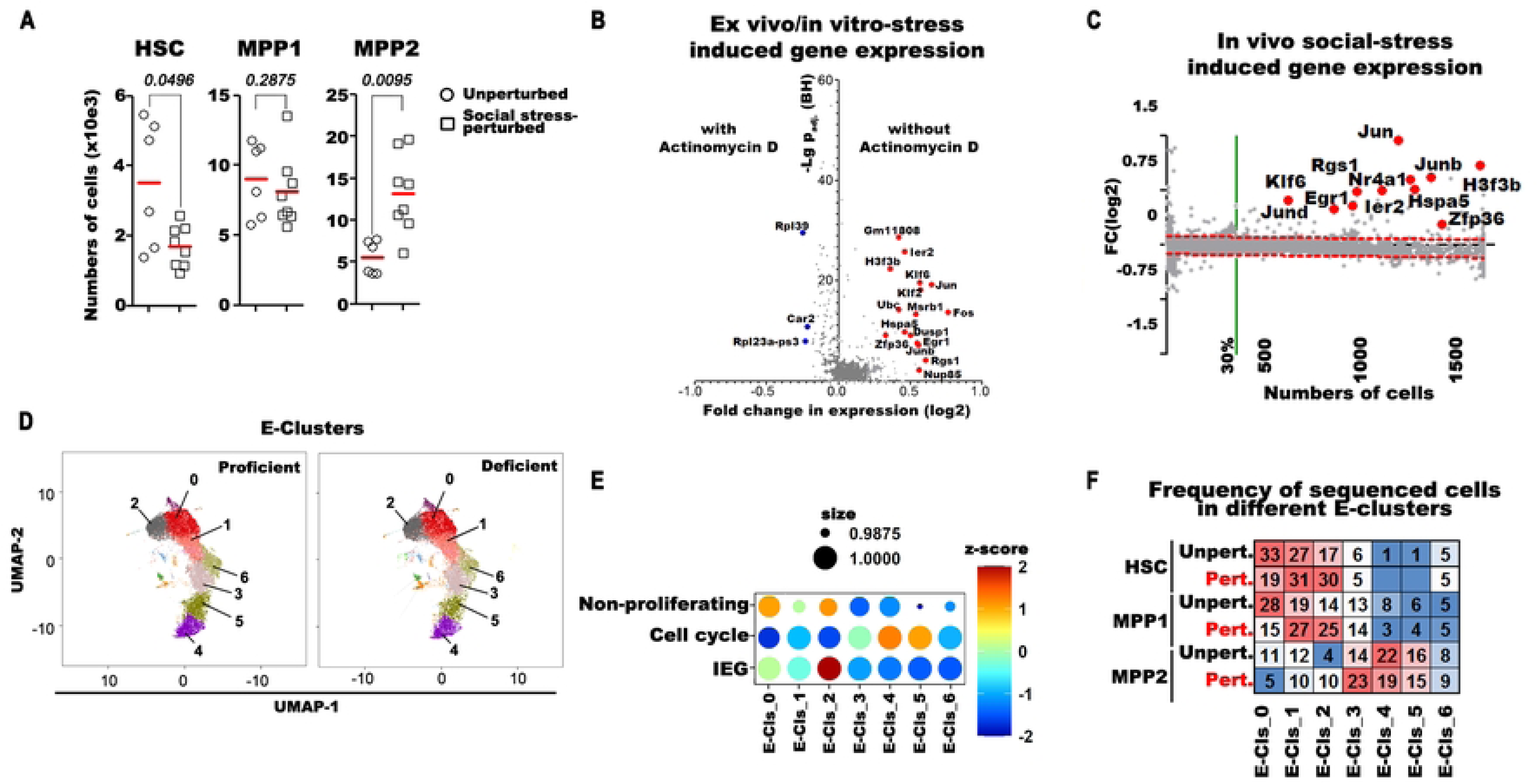
The effect of in vivo social stress- and in vitro/ex vivo stress-perturbation on hematopoietic cells in bone marrow. (**A**) The analysis of flow-cytometry measurements on bone marrow derived hematopoietic stem cells (HSC) multipotent progenitor (MPP) 1 and 2 cells before and after short term in vivo social stress-perturbation. Single cell suspensions from 2 tibia and 2 femurs/mice of unperturbed (open circle) or in vivo social stress perturbed (open square) C57B6/J^TgVav(iCre)+^ mice were stained for flow cytometry, measured and the total numbers of cells/mice were plotted. Red lines indicate the mean values. Paired t-test was used to evaluate statistical significance, exact *p* values are given above the comparing lines. (**B**) The pool of HSC+MPP1 of 2 Actinomycin D treated and 2 non-treated mice and sorted for separate single-cell RNAseq analysis. After data aggregation, 500 genes were selected which were expressed in more than 10% of the cells and had more than 5UMIs. Volcano plot was used to shows statistical significance (logP_adj._BH value) versus magnitude of change (log2 fold change) of expressed genes. Red and blue dots indicate selected genes, have significantly different expression in the presence (blue) or absence (red) of Actinomycin D. (**C**) Genes with higher expression after short-term in vivo social stress perturbation (red dots and gene symbols are selected genes) in HSC are presented by comparative differential expression analysis of unperturbed versus perturbed cells. Differentially expressed genes are above the significance limit. The log2 fold-change expression values were plotted against the numbers of unperturbed cells express the gene. (**D**) The transcriptomes of single HSC, MPP1 and MPP2 cells were analyzed by droplet-based scRNA sequencing. The clustering of early (E) hematopoietic compartment are referred as E-Clusters 0-6. (**E**) Expression of gene-set modules (“Non-proliferating”, “Cell-cycle” and immediate early genes: “IEG”) and coupled frequencies of cells expressing the gene-set in the different clusters are presented as Bubble-plots. (**F**) The distribution of sorted unperturbed (unpert.) and short-time in vivo social stress-perturbed (pert.) HSC, MPP1 and MPP2 cells is shown in the aggregated E-clusters. The frequency of a given cell type in E-clusters is colored from highest to lowest respective red to blue.

### Controlled “ex vivo/in vitro” stress induce transcriptional changes of selected genes, among them Immediate Early Genes

“Ex vivo/in vitro” preparations(10, 11) of hematopoietic cells induce changes in mRNA gene expressions by stimulating transcription of selected sets of genes. Actinomycin D (ActD) irreversibly inhibits transcription(25–27). In order to analyze the “in vitro” stress-induced changes in transcription, we conducted comparative single cell transcriptome analysis on pools of HSC and MPP1 cells prepared in presence or absence of ActD (see Materials and Methods). We found more than 70 genes inhibited by ActD in their “in-vitro” stress-induced increased expression, among them a set of Immediate Early Genes (IEG) (**Fig 1B, Supplementary Table 1**). To avoid this “in vitro” stress-induced transcription, thus, to be able to monitor transcriptional changes induced in HSC and MPP either by “in vivo” stress or, later, by miR-221/222-deficiency, we prepared all cells used in subsequent single cell transcriptome analyses of this study in the presence of ActD.

### Perturbation by “in vivo” social stress induces increased transcription of selected genes, among them IEG in HSC, MPP1 and MPP2 cells

Next, we focused our transcriptome analyses on the earliest progenitors HSC, MPP1 and MPP2 to monitor early changes in gene expression, imposed by the change from the steady-state to the social stress-perturbed state of HSC, MPP1 and MPP2 cells. All sequencing data were analyzed at a similar depth, displaying comparable number of genes and mRNAs per cell, allowing quantitative differential gene expression analyses.

We defined differentially expressed genes as expressed by (i) more than 30% of all unperturbed cells, (ii) with a significantly higher log2 fold-change between unperturbed and “in vivo” social stress perturbed cells, when adjusted to the 99%-prediction band of a linear regression model. Genes above the noise level (red-dashed lines in **Fig 1C** **and Supplementary Fig 2)**, are considered as significantly higher expressed in social stress perturbed cells.

Comparison of gene expression levels in unperturbed and in socially stressed, perturbed HSC (**Fig 1C**), MPP1 or MPP2 (**Supplementary Fig 2**) detected a perturbation-dependent general increase, and an even more pronounced increase in the levels of transcription of a small set of 60-70 genes in HSC, MPP1 and MPP2 cells (**Fig 1C**). This suggests, that perturbation of hematopoiesis by “in vivo” social and by “in vitro” stress might share, at least in parts, the use of IEG-controlled activation.

### Comparison of whole transcriptomes of unperturbed with socially perturbed HSC, MPP1 and MPP2 detects differences in clusters of gene expressions

In primary cluster analysis of the transcriptomes of unperturbed and socially perturbed HSC, MPP1 and MPP2, seven E-clusters (E for early hematopoiesis) were identified (**Fig 1D**). E-0 represents non-proliferating cells (mainly from HSC), E-1 directed towards cell-cycle, and E-2 activated HSC (**Fig 1E**). By location on the UMAP, E-2 is not directed towards proliferating clusters, but its position may indicate an alternative activation. E-3-6 contain MPP1 and MPP2 expressing G1/S and G2/M cell-cycle genes (**Fig 1E; Supplementary Fig 3A; Supplementary/online Table1**).

Compared with unperturbed cells, social stress-perturbed cells in E-clusters were 14% less in non-proliferating E-0, 4% more in proliferation-activated E-1, 9% more in E-3and 13% more in E-2 (**Fig 1F**).

The E-2 cluster contained IEG signatures (**Fig 1E**). Since its position within the UMAP landscape of clusters of transcriptionally relates cells is outside the G1/S/G2-cell cycle-active clusters E1 to E6, they may be differently activated. (**Fig 1D,E**). These results show that early progenitors are activated from quiescence by the stimulatory activity of short-term perturbation.

### The miR-221/222 gene cluster is expressed and can be deleted in all HSC and MPPs

The miR-221/222 gene cluster, located on the Y-chromosome and is expressed in HSC, in activated multipotent progenitors MPP1 and in proliferating MPP2, in myeloid-lymphoid-directed MPP3 and MPP4, and in CLP, granulocytes and myeloid cells. It is turned off in natural killer (NK), thymocytes and T lymphocytes, and preB and B lymphocytes (**Supplementary Fig 4A**).

We determined, how many single HSC, MPP1 and MPP2 express how many miR-221 and miR-222 molecules per cell, by a standard curve-based, multiplexed stem-loop RT-TaqMan qPCR assay. We found that every tested HSC, MPP1 and MPP2 cell expressed between 7000 and 8000 copies of miR-221. Expression of miR-222, again tested in every cell, was lower and more variable, between 1000 and 5000 copies per cell (**Fig 2A**). The level of miR-221 expression strongly correlated with miR-222 expression (**Fig 2B**).

**Fig 2:**
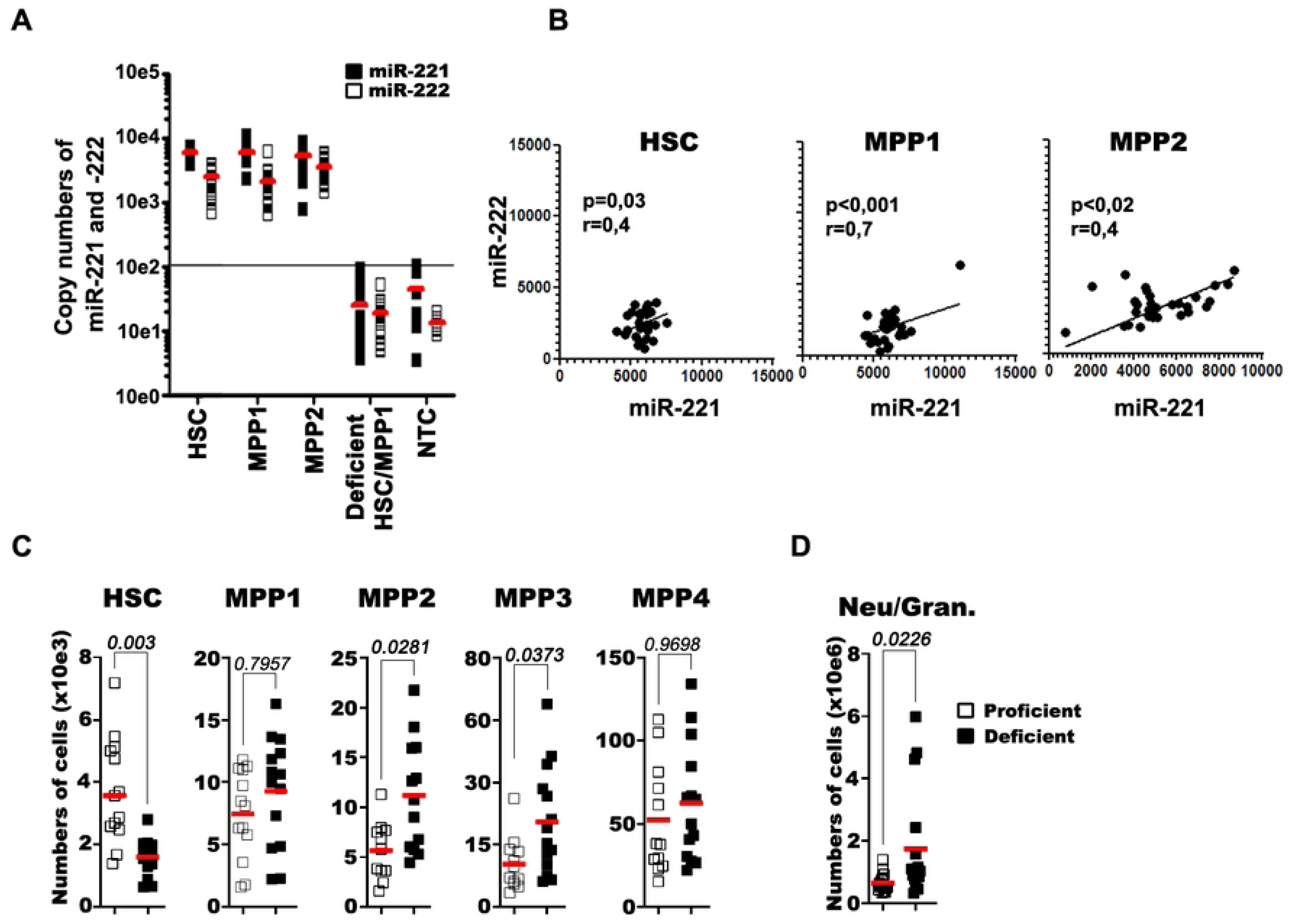
Expression analysis of miR-221/222 in single cells and the effect of miRNA deficiency on hematopoietic stem cell populations in bone marrow. (**A**) Copy numbers of miR-221 (closed square) and miR-222 (open square) measured in single HSC, MPP1 and MPP2 of miRNA-proficient and in pools of HSC+MPP1 single cells of miRNA-deficient mice (n_cell_= 36 cells from 3 mice, n_NTC_= 12 wells). NTC: no-template control. The highest value of NTC is marked as straight line, indicating the detection limit. (**B**) Correlation analysis of miR-221 and miR-222 copy numbers in proficient HSC, MPP1 and MPP2 cells (**C, D**) The analysis of flow-cytometry measurements on bone marrow and spleen derived cells. (**C**) Single cell suspensions from 2 tibia and 2 femurs/mice or (**D**) from spleens of miR-221/222-proficient (open square) or deficient (closed square) mice were stained for flow cytometry, measured and the total numbers of cells/mice were plotted. Red lines indicate the mean values. Paired t-test was used to evaluate statistical significance, exact *p* values are given above the comparing lines.

To study functions of miR-221/222 during hematopoiesis, we generated mice, in which the one miR-221/222 cluster on the Y-chromosome in male mice could be deleted in all hematopoietic cells. Vav^i-cre^, active in all hematopoietic cells including HSCs, is commonly used to drive Cre recombinase expression in hematopoietic cells(4, 42). Thus, Vav-Cre mice were crossed with miR-221/222^fl/fl^ mice to generate male F1 progeny, where the miR-221/222 cluster was expected to be deleted on the one Y-chromosome. All mouse strains had been backcrossed on C57BL/6J for at least 8 generations to ascertain genetic homogeneity, except for the miR-221/222 cluster in all offsprings. Bulk-sorted BM-lineage (lin)^-^c-kit^+^Sca1^+^ (LSK), and multipotent progenitor cells (LSK CD150^-^CD48^+^) (**Supplementary Fig 4B**), and single LSK CD150^+^CD48^-^ (pool of HSC and MPP1) cells of miR-221/222^fl/y- Tg(Vav1-icre)^ mice were tested for quantitative miR-221/222 expression at single cell level. Indeed, our analyses show high efficiency of deletion of the miR-221/222 cluster in all tested cell populations. (**Fig 2A, Supplementary Fig 4B**).

### MiR-221/222-deficiency decreases HSC and increases MPP in unperturbed hematopoiesis

In BM of miR-221/222-deficient mice at steady state of unperturbed hematopoiesis, we found numbers of HSC reduced approximately threefold, whereas MPP2 were increased. (**Fig 2C**). The extent of these changes in the numbers of HSC, MPP1 and MPP2 are similar to those seen in socially perturbed HSC, MPP1 and MPP2. This could suggest, that miR-221/222-deficiency and social stress might use, at least in parts, the same mechanisms to regulate pool sizes of early progenitors.

Numbers of MPP1, MPP4, CLP, CMP, megakaryocyte-erythroid (MEP) and granulocyte-myelocyte (GMP) progenitors and peripheral CD4^+^, CD8^+^ T cells and CD19^+^ B cells in spleen were not changed (**Fig 2C, Supplementary Fig 1A,B**). Interestingly, in spleen the numbers of granulocytes were 6-fold elevated in miR-221/222-deficient mice (**Fig 2D**). These results suggest that expression of miR-221/222 maintains pool sizes of HSC and MPPs and prevents an increased granulopoiesis.

### MiR-221/222 targets *Fos* to protect the size of the unperturbed HSC pool

Next, we compared transcriptomes of deficient with proficient cells to detect microRNA-sensitive genes. We defined these genes as expressed by (i) more than 30% of all miR-221/222-proficien cells, (ii) with a significantly higher log2 fold-change between proficient and miR-221/222-deficient cells, when adjusted to the 99%-prediction band of a linear regression model. Genes above the noise level (red-dashed lines in **Fig 3A,E,F**), are considered as significantly expressed in miR-221/222-deficient cells.

**Fig 3:**
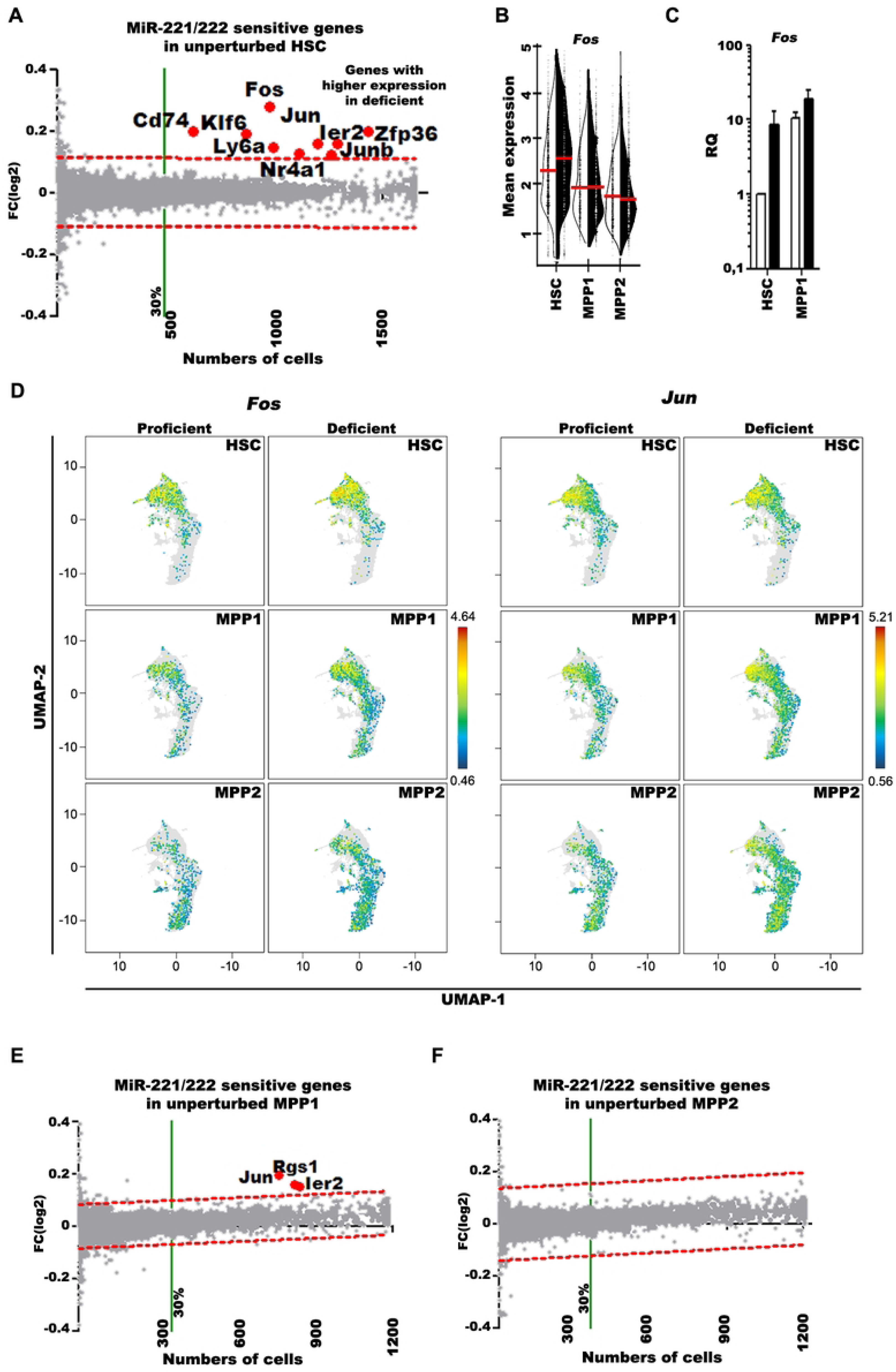
MiR-221/222 sensitive genes in unperturbed HSC, MPP1 and MPP2 cells. (**A**) Comparative differential expression analysis between miR-221/222-deficient and proficient HSC, (**E**) MPP1 and (**F**) MPP2 cells. The log2 fold-change expression values are plotted against the numbers of proficient cells expressing the gene. After calculating the linear regression curve, the 99% confidence interval (CI) band (dashed red lines) determined the non-differentially (gray dots) and differentially (red dots and gene symbols) expressed genes. Significantly higher expressed genes in miR-221/222-deficient cells are above the 99% CI limit and are expressed by more than 30% of all cells (green line). These genes are called “differentially expressed genes”. (**B**) Single cell expression of *Fos* in HSC, MPP1 and MPP2 cells. The difference in mean expression level of the indicated populations between proficient (open half violin) and deficient (filled half violin) cells are presented (**C**) The relative quantity (RQ) of *Fos* mRNA in bulk sorted miR-221/222-proficient (open bars) and -deficient (filled bar) HSC and MPP1 cells measured by RT-qPCR. RQ was calculated on the basis of *Hprt* expression in miR-221/222-proficient HSC. (**D**) The differential expression of AP-1 components *Fos* and *Jun* presented on UMAP of unperturbed miR-221/222 proficient and deficient HSC, MPP1 and MPP2 cells.

At steady state of unperturbed hematopoiesis, we detected nine such differentially expressed genes (**Fig 3A,B**). Among them, *Fos* was the only miR-221/222-seed-sequence-containing target gene (DIANA-TarBase v8(48), TargetScan Release 8.0(49), miRDB(50)). We validated this increased *Fos* expression with qPCR analyses (**Fig 3C**), in which *Fos* was found to be expressed 8.5-fold higher in deficient HSC. Six of the other 8 genes (*Jun*, *JunB*, *Nr4a1*, *Ier2*, *Zfp36* and *Klf6*) are, like *Fos* IEG(43, 51) (**Fig 3A,D**).

Next, we performed the same analyses with MPP1 and MPP2. In unperturbed MPP1, *Fos*, *JunB*, *Nr4a1*, *Cd74*, *Ly6a*, *Zfp36* and *Klf6* were no longer detectable, and *Rgs1* appeared (**Fig 3D,E**). In MPP2, no differentially expressed genes remained detectable (**Fig 3F**). This suggests that in unperturbed hematopoiesis upregulated mRNA expression of *Fos*, *Jun*, and five other IEG transcription factors act primarily in HSC, causing a decrease of HSC and an increase of MPPs and of granulocytes.

### Shared IEG expression upregulated by social stress-perturbation, “in vitro”-perturbation and miR-221/222-deficiency

Next, we compared the differentially upregulated genes in transcriptomes of 1) “in vitro” perturbed, 2) of social stress-perturbed and of 3) experimentally unperturbed, miR-221/222-deficiency-influenced HSC and MPP1. Six IEGs – *Fos*, *Jun*, *JunB*, *Ier2*, *Klf6* and *Zfp36* – were found up-regulated, by all three stressors (**Fig 4A**). This suggests, that HSC and MPP1 use IEG responses to control pool sizes of HSC and their differentiation to granulocytes. The results of our experiments indicate, that this pathway is controlled by the miR-221/222 gene cluster.

**Fig 4:**
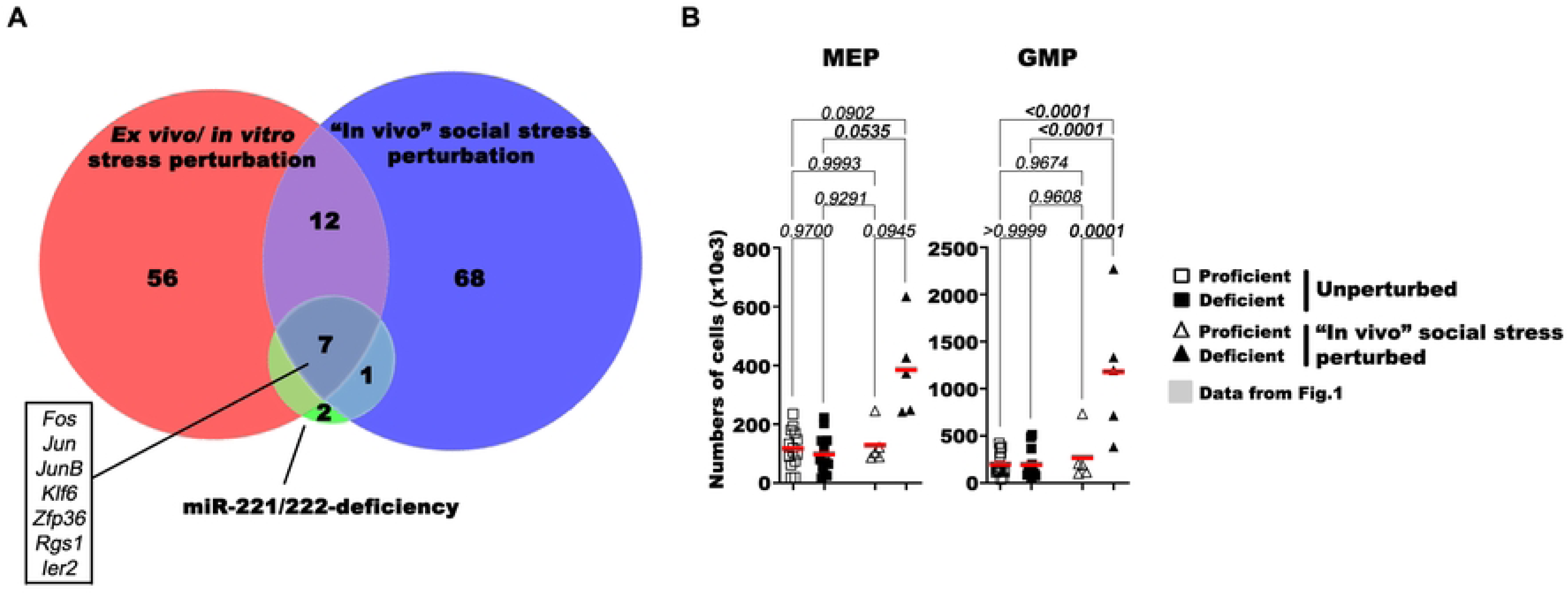
Shared gene expression program in ex vivo/in vitro stress-perturbed, in vivo social stress-perturbed and miR-221/222-deficiency-induced HSC and MPP1 cells. (**A**) The ex vivo/in vitro stress-perturbed, in vivo social stress-perturbed and miR-221/222-deficiency-induced genes were determined by differential gene expression analyses of scRNA-seq data and the plotted. The area-proportional Venn-diagram shows the commonality between ex-vivo perturbation sensitive, short-term social stress perturbation sensitive and miR-221/222-deficency sensitive genes. (**B**) Analysis of flow-cytometry measurements on unperturbed and social-stress perturbed myeloid-erythroid progenitors (MEP) and granuloid-myeloid progenitors (GMP). Single cell suspensions of tibia and femurs of unperturbed miR-221/222-proficient (open squares) and miR-221/222-deficient (closed squares) mice or perturbed miR-221/222-proficient (open triangles) or miR-221/222-deficient (closed triangle) mice were prepared, analyzed with flow-cytometry and the numbers of cells were plotted. Red lines indicate the mean values. One-way ANOVA with Tukey post-test was used to evaluate statistical significance, exact *p* values are given above the comparing lines.

### Controlled perturbation by social stress of miR-221/222-deficient BM leads to increases in numbers of MEP and GMP granulocyte progenitors

In unperturbed, miR-221/222-deficient mice increased numbers of granulocytes had been observed as a possible involvement of this miRNA cluster in enhanced granulopoiesis (**Fig 2D**). In support of this notion, short social stress (20 hours) was sufficient to induce increases in the numbers of MEP and GMP (in average 4-fold each) in miR-221/222-deficient, but not in miR-221/222-proficient BM (**Fig 4B**). The number of CLP and CMP was not different. These rapid changes in granulocyte progenitor compartment sizes suggest, that miR-221/222-deficient, stressed HSC and MPPs need only a short period of perturbation, in which proliferation and differentiation increase granulocyte-biased, but not lymphoid-directed hematopoiesis, detectable in increased numbers of MEP and GMP. By contrast, stress induced changes in HSC and MPP numbers are not further altered by miRNA-deficiency. Due probably to the short period of activation, the numbers of mature granulocytes in the periphery are not (yet) further increased. We conclude from these results, that miR-221/222 safeguards myeloid-lymphoid multipotency of HSC and MPP also in stressed hematopoiesis.

### Transcriptional landscapes of hematopoietic progenitor cells detect miR-221/222-deficiency-induced granulopoiesis

In order to construct comprehensive transcriptional landscapes of hematopoietic progenitors in BM at unperturbed, steady-state hematopoiesis, with trajectories of gene expressions during development to different lymphoid, myeloid and erythroid cell lineages, and to detect possible changes in granulocyte-directed trajectories, we conducted separate transcriptome analyses for HSC, MPP1 and MPP2, including also more mature BM progenitors, i.e. MPP3, MPP4, CLP and lin^-^c-kit^+^Sca1^-^ or lin^-^c-kit^-^Sca1^-^ cells from miR-221/222-proficient and deficient BM. Cells were clustered (T-clusters for “total”) according to their transcriptomes by Uniform Manifold Approximation and Projection for Dimension Reduction (UMAP)(52) (**Fig 5A** and Supplementary methods).

**Fig 5:**
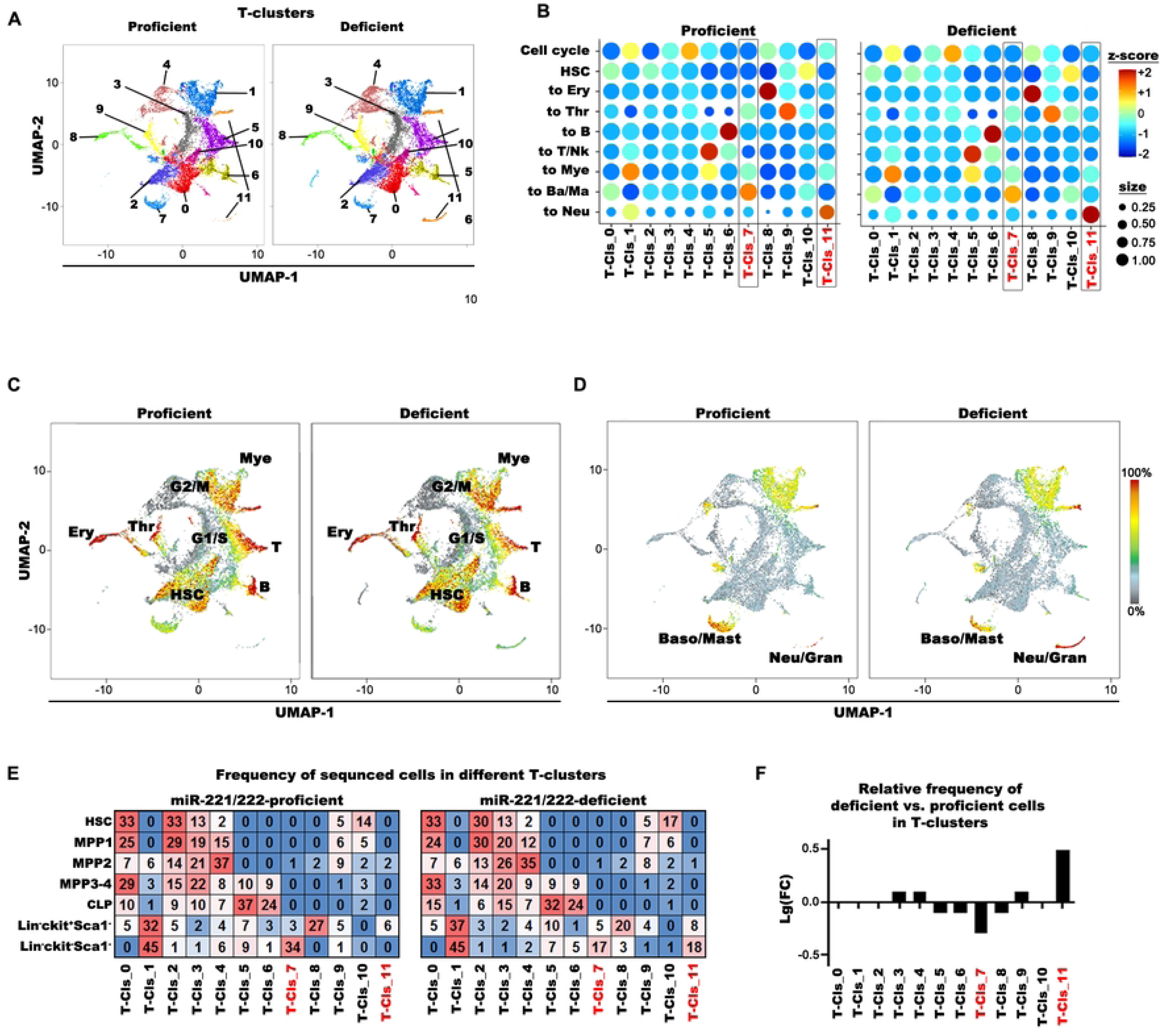
Single cell transcriptome analysis of unperturbed miR-221/222-proficient and deficient lineage negative bone marrow compartment. (**A**) Cluster analysis on the aggregated Uniform Manifest Approximation Projection (UMAP) plots of miR-221/222-proficient and deficient HSC, MPP1, MPP2, MPP3-4 pool, CLP, lin^-^ ckit^+^Sca1^-^ and lin^-^ckit^-^Sca1^-^ populations (total, T clusters) after single cell-transcriptome sequencing. Proficient and deficient cells are separately plotted. (**B**) The bubble-plots shows the expression of gene-set modules (z-score) and coupled frequencies of cells (bubble size) expressing the gene-set in the different T-clusters. MiR-221/222-proficient and deficient samples were separately plotted. (**C, D**) Major hematopoietic differentiation pathways are presented on the UMAP as characteristic gene-set modules (**Supplementary Table 1**) by the Log2 summary expression of the module genes. (**C**) Cells with characteristic expression pattern for Mye (to Myeloid), T (to T cells), B (to B cells), Ery (to erythrocytes), Thr (to thrombocytes) are shown on a composite picture for proficient and deficient cells and (**D**) for Ba/Ma (to basophil/mast cells) and Neu/Gran. (to neutrophils/granulocytes). (**E**) Frequencies of sorted miR-221/222 proficient and deficient HSC, MPP1, MPP2, MPP3-4 pool, CLP, lin^-^ckit^+^Sca1^-^ and lin^-^ckit^-^Sca1^-^ cells in different T-clusters. The frequency of a given cell type in T-clusters is colored from highest to lowest respective red to blue. (**F**) Relative frequencies of cells in different T-clusters are presented on Log10 scale (frequency of deficient cells relative to the frequency of proficient cells).

We selected sets of genes to characterize expression programs for HSC, cell-cycle, erythroid, megakaryocytic, lymphoid, myeloid and granulocyte differentiation and assigned these T-clusters (**Fig 5B-E; Supplementary Fig 3B** and **Supplementary Table 1**). These analyses are in agreement with earlier publications of other laboratories(1–8, 53).

Only two T-clusters were different in proficient and deficient cells. T-7 (basophil/mast cell) contained -1.9-fold less, T-11 (granulocytes) 2.9-fold more cells in miR-221/222-deficient mice (**Fig 5E,F**). In fact, the granulocyte-related genes were only present in T-11 of miR-221/222-deficient, but not of proficient granulocytes (**Fig 5B,D**, lower left- and right-hand corners).

These results support our findings (**Fig 4B**) of increased numbers of granulocyte precursors in BM and granulocytes in the spleen miR-221/222-deficient mice. They suggest, that steady-state hematopoiesis of miR-221/222-deficient BM progenitor cells increase granulopoiesis even in an experimentally unperturbed state.

### In perturbed HSC, MPP1 and MPP2 cells miR-221/222-deficiency selectively increases transcription of genes encoding heat shock proteins, G-protein-mediated tubulin and chromatin remodeling activities

Finally, we compared transcriptomes of miR-221/222-deficient with proficient cells to detect microRNA-sensitive genes after “in vivo” social stress perturbation. We defined differentially expressed genes as expressed by (i) more than 30% of all in vivo social stress perturbed miR-221/222-proficient cells, (ii) with a significantly higher log2 fold-change between in-vivo social stress perturbed miR-221/222-proficient and deficient cells, when adjusted to the 99%-prediction band of a linear regression model. Genes above the noise level (red-dashed lines in **Fig 6A-C**), are considered as significantly expressed in in-vivo social stress perturbed miR-221/222-deficient cells.

**Fig 6:**
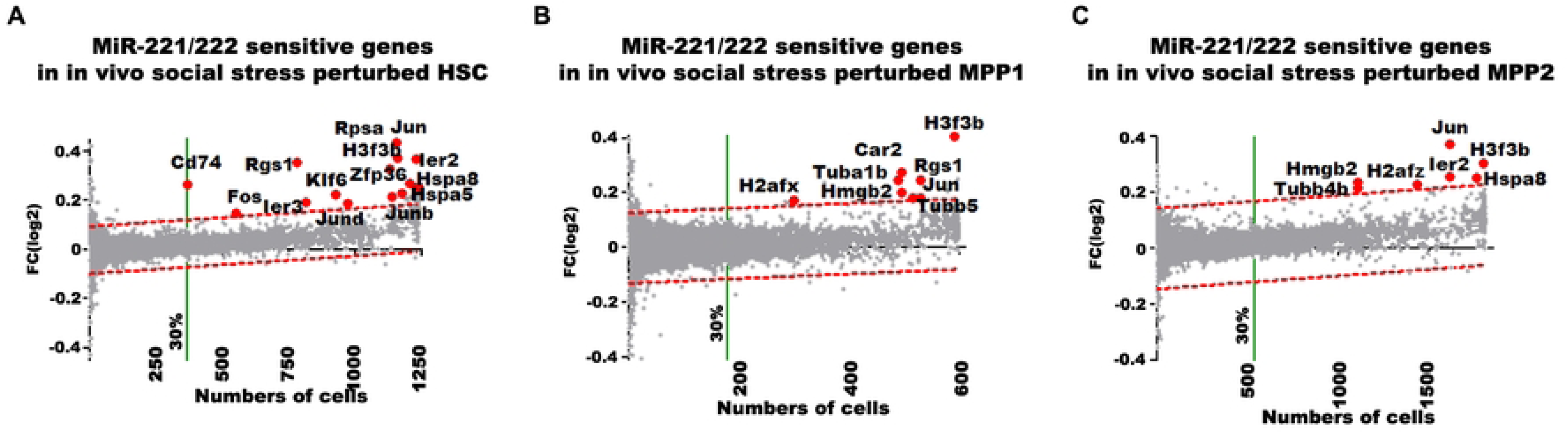
Analysis of in vivo social stress-perturbation sensitive genes in miR-221/222-deficient HSC, MPP1 and MPP2 cells. Genes with higher expression after short-term perturbation (red dots and gene symbols are selected genes) in miR-221/222-deficient (**A**) HSC (**B**) MPP1 and (**C**) MPP2 are presented by comparative differential expression analysis of unperturbed versus perturbed cells. Differentially expressed genes are above the significance limit. The log2 fold-change expression values were plotted against the numbers of unperturbed cells express the gene.

Beside upregulated IEG expression, social stress-induction combined with miR-221/222-deficiency selectively upregulated three groups of functionally connected genes in HSC, MPP1 and/or MPP2 (**Fig 6A-C**):

1. Heat shock protein (HSP70) genes *Hspa5* and *Hspa8*, with possibilities to activate the unfolded protein response (UPR)(54).
2. G-protein signaling (*Rgs1*) and its tubulin components (*Tubb1b*, *4b* and *5*) in cytoskeleton and microtubule assembly, spindle formation and mitotic cell cycle control(55). Beyond the known activities of Rgs1 in hypoxia(56) and in SDF-1/CXCR4-induced migration of HSC(57), these genes are expected to contribute to increased erythroid-myeloid differentiation.
3. Replication-independent, histone-associated chromatin remodelers (*H3f3b*, *Hmgb2*, *H2afx*, *H2afz*). Since *H3f3b* is known to induce erythroid differentiation(58), miR-221/222-dependent upregulation of *H3f3b* would favor erythroid-myeloid progenitor activation, as seen in our analyses.

We expect, that the combined actions of all of these up-regulated genes could contribute to the activation of HSC from quiescence and to induce selectively increased granulopoiesis(53, 59–62)

### MiR-221/222-deficient HSC fail to reconstitute recipients in serial transplantations

In transplantations with the aim to reconstitute the hematopoietic cells of a lethally irradiated host, CD34^-^ HSCs have been found to contain long-term, fully repopulating HSC, while CD34^+^ MPP1 repopulate all lineages, but have reduced capacities to reconstitute long-term repopulating HSC and early progenitors(1, 4, 5). Serial transplantations of HSC can be expected to impose “in vitro” stress on HSC during their preparation for transplantations.

In order to test repopulation capacities of miR-221/222-proficient and of deficient HSC, we transplanted 100 donor CD45.2^+^ either proficient or deficient HSC into lethally irradiated CD45.1^+^ mice, together with 10^6^ non-irradiated CD45.1^+^ BM carrier cells known to contain 100 proficient HSC – conditions, which are expected to lead to 50% CD45.1-50% CD45.2 chimerism. In serial transplantations we then sorted HSC from the transplanted mice and re-transplanted 100 HSC under the same conditions.

In blood, the chimerism in total CD45^+^ hematopoietic cells was found to be close to these expected values after the first (∼40%) and second (∼55%) transplantations of miR-221/222-proficient HSC (**Fig 7A**). With miR-221/222-deficient HSC, this chimerism was significantly lower 4 months after the first (∼24%), and much lower 4 months after the second (2,5%) transplantations, suggesting, that miR-221/222-deficiency exhausts the capacity of HSC for hematopoietic reconstitution.

**Fig 7:**
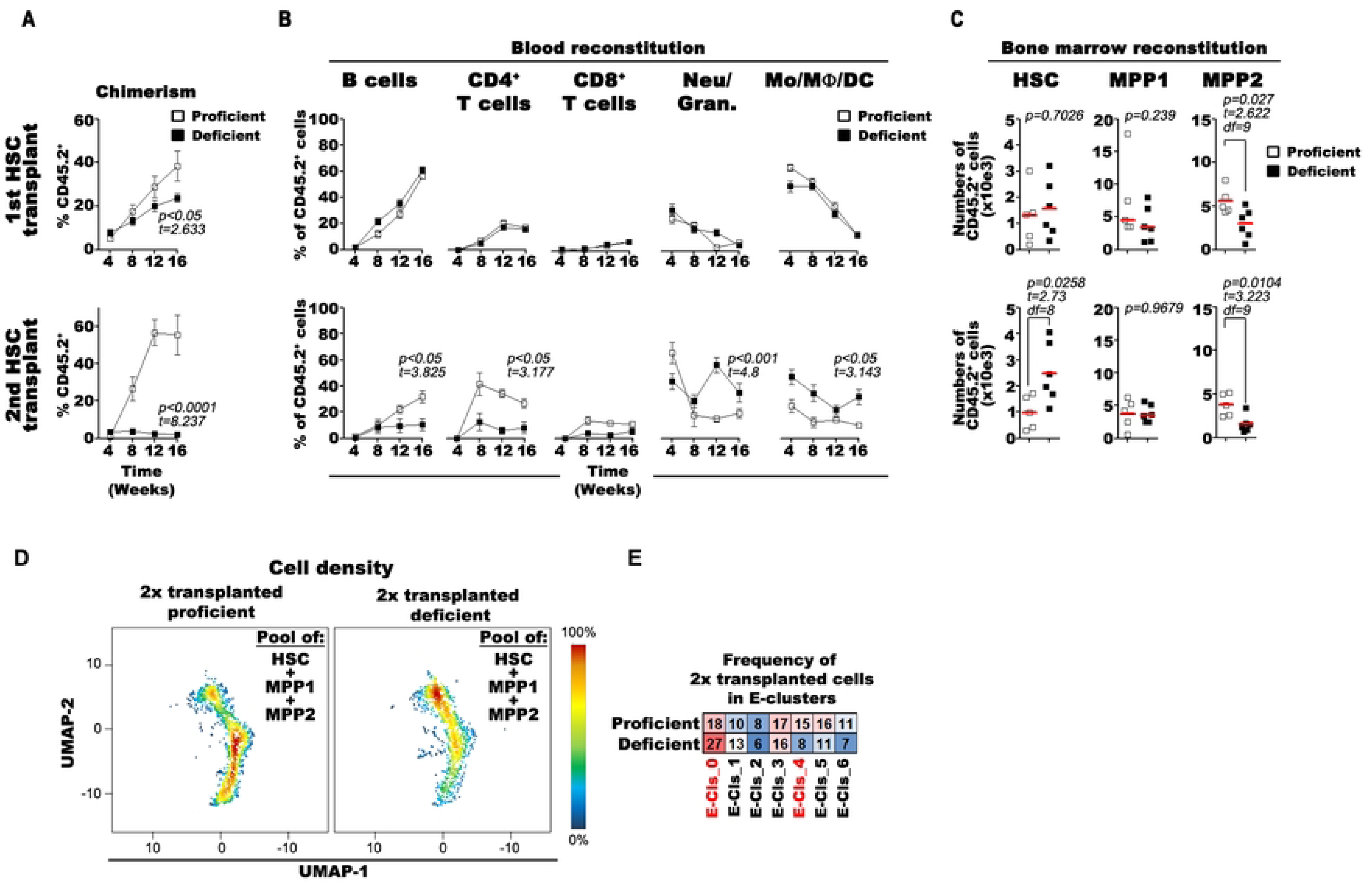
MiR-221/222 cluster safeguards the multipotency of HSC. (**A**) Proportion of donor-derived blood cells after the first (upper) and second (lower) transplantation of 100 miR-221/222 proficient (open squares) and deficient (closed squares) HSC. (**B**) Frequencies of B cells, CD4^+^/CD8^+^ T cells, CD11b^+^Gr1^+^ neutrophils/granulocytes (Neu/Gran.) and CD11b^+^Gr1^-^ monocytes/macrophages/dendritic cells (Mo/MF/DC) of donor derived blood cells after the first (upper line) or second (lower line) transplantation. (**C**) Numbers of donor-derived HSC, MPP1 and MPP2 cells 16 weeks after the first (upper line) or the second (lower line) miR-221/222 proficient (open squares) and deficient (closed squares) HSC transplantation. Red lines indicate the mean values. (**A-C**) Data from 5 miR-221/222-proficient and from 6 deficient mice are presented as (**A,B**) mean values or (**C**) individually. Paired t-test was used to calculate significant differences, exact *p* values are given above the comparing lines. (**D**) Density plots of pooled miR-221/222 –proficient and deficient HSC, MPP1 and MPP2 cells after serial transplantation are visualized on the aggregated UMAP and (**E**) the distribution of cells in the different E-clusters (see clusters on (Fig 1D)). The frequency of a given cell type in E-clusters is colored from highest to lowest respective red to blue.

Despite this reduced peripheral chimerism in deficient versus proficient T (15% vs 35%) and B cells (10% vs. 30%), no differences in reconstitution were observed between proficient and deficient HSCs in BM 4 months after the second transplantation. Furthermore, secondary recipients of proficient HSC had 30% donor-derived myeloid cells and granulocytes, while recipients of deficient HSC contained 65%. This suggests, that twice-transplanted miR-221/222-deficient HSC continued to develop granulocytes, but had become deficient in their capacities to develop lymphoid cells (**Fig 7B**).

In BM, serial transplantations of proficient and deficient HSC generated different sizes of repopulated HSC and MPP2 pools (**Fig 7C**). We found an equal chimerism of proficient as well as deficient CD45.2^+^ HSC after the first transplantation, indicating comparable, adequate homing to the bone marrow of both types of HSC. Four months after secondary transplantation, proficient HSC established normal HSC levels. Surprisingly, levels of twice transplanted deficient HSC levels were even 2 to 3-fold higher (**Fig 7C**). Numbers of MPP1 were not different, while MPP2 were reduced already after the first, and even more after the second transplantation of deficient HSC (**Fig 7C**). These data indicate that twice transplanted miR-221/222-deficient HSC retain their capacity to home to and repopulate their niches in BM, but lose parts of their capacities of hematopoietic differentiation. Thus, the repeated “in vitro” stress during transplantation might favor the preservation, or even enhancement of granulocyte differentiation capacities of miR-221/222-deficient HSC, but it also extinguishes their lymphoid differentiation capabilities.

### Serial transplantations of miR-221/222-deficient HSC accumulate cell cycle-inactive HSC and deplete proliferating MPPs

Finally, we also compared twice serially transplanted miR-221/222-deficient HSC and MPPs, which had lost their multipotency, but had retained their BM-homing-capacity (**Fig 7A-C**) with proficient, twice-transplanted progenitors. MiR-221/222-deficient cells had increased numbers of E-0 (non-proliferating HSC), but decreased numbers of E-4-6 (cell-cycle-active) (**Fig 7D,E**). This suggests that serial transplantation of miR-221/222-deficient HSC accumulates proliferation-inactive HSC, and depletes cell-cycle-active MPPs. These deficient HSC are reminiscent of differentiation-inactive HSC clones described by Pei et al.(8). Again, we conclude from these results, that miR-221/222 safeguards the multipotency of HSC.

## DISCUSSION

Most central and peripheral hematopoietic cell compartments turn over throughout life, with half-lives between a few days as a few weeks. By contrast, most HSC are generated once during embryonic development of the bone and its marrow, to remain long-lived, quiescent cells throughout life(1–9). A smaller part of these HSCs and more differentiated progenitors serve as life-long sources for this continuous regeneration. In mouse BM a few thousand LSK CD150^+^ CD48^-^ CD34^-^ HSC have the capacity to survive as quiescent cells without dividing for years, even for life. When transplanted into recipient mice, a single HSC can repopulate the HSC compartment, all progenitor compartments and all the mature hematopoietic lineages in normal numbers. Hence, HSC can find their niches in bone marrow, can fill these niches by symmetric cell divisions, and can initiate multi-lineage differentiation to mature erythroid, megakaryocytic, myeloid and lymphoid cells(1–9).

Perturbations of hematopoiesis by bacterial(15) or viral infections(13), by social stress(12), or by transplantation(1, 4), mediated by interferon-α(18) or interferon-γ(19), by poly-I:C dsRNA(15) or by G-CSF(20) have all been found to restrict the multi-lineage differentiation capacities of HSC and favor differentiation-restricted HSC, leading e.g to emergency granulopoiesis(21). Such restrictions have been seen to become more prominent with aging(22, 24). Repeated, long-term perturbations of HSC during life, e.g. by infections(13), by repeated social stress(12) or during serial BM transplantations, as done in our experiments, all can contribute to HSC aging(36). Young HSC maintain an apparently more unperturbed balance between quiescence, activation and multipotent lymphoid-myeloid differentiation. With age, less HSC appear quiescent, more are activated, less are lymphoid differentiation-directed and more are myeloid differentiation-directed (1–9).

Several miRNAs have been identified, which influence the regulations of these different states of HSC(36). Several of them (miR-21, miR-22, miR-99, miR-125a/b and miR-155, reviewed in Luinenburg et al.(36) are target genes in the mTOR pathway(37). Our results identify the miR-221/222 cluster as a set of new miRNA genes, which regulate HSC by a new pathway of cellular activation - via Fos-AP-1-IEG – to protect their quiescence and multipotent lymphoid-myeloid differentiation. Interestingly, while relatively little is known about the actions of Fos in HSC, expression of Fos and of other IEGs is used since a long time as a marker of cell activation by individual neurons(63).

In our present work we have studied the role of the microRNA cluster 221/222 which is expressed at high levels in hematopoietic precursors including HSC, during steady state and stressed hematopoiesis. While we find the homing capacities of HSC unaltered by miR-221/222, we show that the expression of miR-221/222 in HSC is indispensable for safeguarding the HSC cell pool from perturbation-induced depletion, granulopoiesis-poised differentiation and the loss of lymphoid reconstitution potential. Thus, we have identified miR-221/222 cluster as a new key regulator of HSC identity. We have found, that all HSC and MPP1 express miR-221/222(35, 64) (**Fig 1B**). Furthermore, we found, that Vav-iCre deleted the miR-221/222 in all HSC and MPP1 cells. Nevertheless, not all HSC and MPPs, but only a larger part of them is activated from HSC to MPPs, when deficiency is induced.

We have identified Fos as the only direct target gene containing miR-221/222-seed sequences to be differentially expressed in miRNA-deficient HSC although the bioinformatical tools predict many other target genes which we found expressed, suggesting, that most of the predicted and validated targets are not susceptible to miR-221/222 action in these cells. Again, even with Fos, not all unperturbed HSC, or perturbed HSC show these changes (**Fig 1A** and **2C**). These restrictions in the action of the miR-221/222 cluster remain to be investigated in greater detail.

It is important to realize, that we would not have been able to discover, that mRNAs of the immediate early genes Fos, Jun, Nr4a1, Ier2 and Zfp36 are differentially upregulated in miR-221/222-deficient HSC, if we would not have performed the “ex vivo” preparation of HSC, MPP1 and MPP2 from bone marrow in the presence of ActD, which inhibits RNA synthesis irreversibly(26) and preserves the transcriptional *in vivo* state of the cells. In the absence of ActD, we found HSC and MPPs to be stimulated “in vitro” during cell preparatory steps to transcribe a collection of genes, among them IEGs, which would have obscured the detection of differentially expressed, miR-221/222-sensitive genes, notably of Fos.

In fact, our results suggest, that bone marrow transplantations might be improved, if the “in vitro” stress reactions on HSC could be avoided. Certainly, treatment of HSC with cell-toxic ActD is not an option for improvement, but upregulation of miR-221/222 during HSC preparation prior to transplantation could be, if we were able to control the expression of this miR cluster in HSC.

Most of the published effect of Fos describes an important role in early hematopoiesis, i.e. in the endothelial-to-hematopoietic transition during embryonic establishment of the HSC compartment. Fos (and their family members FosB, Fra-1 and Fra-2) can form hetero-dimeric AP-1 transcription factor complexes with Jun (and their family members JunB and JunD)(43, 65–67). AP-1 regulates a wide collection of cellular responses, such as proliferation, differentiation and apoptosis(43). In early hematopoiesis AP-1 guides development from embryonic stem cells by activating a specific set of genes to develop smooth muscle vascular endothelium. Inhibition of AP-1 shifts the balance between smooth muscle and hematopoietic differentiation in favor of hematopoiesis(68). Furthermore, AP-1 complexes have been found to regulate BMP4-induced hematopoiesis(69). In AP-1 complexes, Fos influences the engraftment of HSC into their niches in bone marrow, a crucial early step in the establishment of niches for their longevity in bone marrow(70).

IEG expression in HSC could be expected to involve, among others, c-kit, CXCR4, MPL, TGFβ− and interferon receptors on the cell surface and various intracellular signaling events leading to JNK-mediated activation of Fos/AP-1 transcription factor. AP-1 has been identified as one of several known transcription regulators, also including SRF, CREB, KROX and NFκB, over-represented in the upstream region of immediate-early genes(44). Our findings, that miR-221/222-deficiency-induced up-regulation of the Fos/AP-1/IEGs, appears sufficient to activate HSC, gives the Fos/AP-1-transcription factors a dominant controlling influence on transcription of only a limited number of IEG, activating HSC from quiescence to cell cycle and to increased granulopoiesis. We show here, that social as well as “in vitro” stress uses these pathways to activate miR-221/222-proficient HSC. Our findings, that Fos/AP-1 upregulation in miR-221/222-deficient HSC, together with indirect upregulation of six IEG genes, suffices to activate non-perturbed miR-221/222-deficient HSC in steady state of hematopoiesis is a strong indication, that Fos and six other IEG genes are, in fact, required for this activation.Klf6, together with Runx1, promotes the transition of neutrophils from bone marrow to blood (71, 72). Our finding, that miR-221/222 selectively upregulates the expression of Klf6 in HSC, that miR-221/222-deficient mice have increased levels of granulocytes in spleen, and that serial transplantation of miR-221/222-deficient HSC leads to the development of myeloid-granulocytic-biased HSC, all may indicate that this promotion to neutrophil development begins in HSC in bone marrow. Nr4a1, an orphan nuclear receptor has been found expressed on myeloid-biased HSC(73). Thus, miR-221/222-deficiency, promoting the upregulated expression of Nr4a1 (**Fig 3A**), might cooperate with Klf6 to condition HSC for myeloid-granulocytic development. Zfp36, an AU-rich RNA-binding protein(74, 75), has been found to suppress hypoxia and cell cycle signaling, activities, which are known influence HSC performances. In gene expression trajectories of neutrophil-granulocyte development from HSC Zfp36 has been identified as a differentiation-determining factor(15, 76, 77). In conclusion, three of the six IEGs selectively up-regulated in unperturbed hematopoiesis by miR-221/222-deficiency-dependent up-regulation of Fos, i.e. of AP-1. Namely Klf6, Nr4a1 and Zfp36 all might cooperate to condition HSC for myeloid-granulocyte-biased hematopoiesis. Since AP-1 can be expected to regulate the expression of many more genes expressed in HSC, it remains to be investigated, how miR-221/222-deficiency focuses its effect in HSC to this selective set of genes, thereby conditioning them for myeloid-granulocytic development(76). Here, we found, that time-controlled perturbation by social stress activated HSC to increased MPP development and to emergency granulopoiesis, helping to strengthen the emergency granulopoiesis induced by miR-221/222-deficiency. In the short time of stress, this support of emergency granulopoiesis was not yet visible by an increase in numbers of mature peripheral granulocytes, but was detectable in the primary environment of bone marrow as dramatic increases in numbers of granulocytic progenitors within less than a day. Stress selectively increases the expression of around one hundred genes, many of them IEGs. Notably, transcription of the direct miR-221/222 target gene Fos/AP1 is also increased, and the increased expression extends beyond HSC into the MPP1 and MPP2 compartments. In the stress-perturbed HSC, MPP1 and/or MPP2 the differentially, miR-221/222-deficiency-dependently up-regulated mRNA expression now includes not only some of the IEGs, but also heat shock protein genes (Hspa5 and Hspa8) with UPR potential, replication-independent histone modifier genes (H3f3b, Hmgb2, H2afx, H2afz) with the potential to alter gene expressions, and the G-coupled protein signaling gene Rgs1 and tubulin genes with the potential to modify cytoskeleton architecture. All of these up-regulated genes are candidates for activities, which activate HSC from quiescence and induce emergency granulopoiesis.

Consequently, miR-221/222 expression is expected to suppress the expression of these differentially expressed genes and, thus, down-regulate stress-induced granulopoiesis at multiple levels. Future studies will have to identify the factors, which regulate the expression of the miR-221/222 gene cluster in early hematopoiesis.

A deeper understanding of the molecular mechanisms, how HSCs regulate engraftment in bone marrow and balance self-renewal vs. differentiation, has important clinical implications for mobilizing defense against infections, for improving bone marrow transplantations, and for fighting cancer of hematopoietic precursor cells. Our findings suggest that expression of the miR-221/222 cluster safeguards long-lived, quiescent HSC from Fos/AP-1-induced activation resulting in the reduction of HSC, the proliferation of MPPs, and their differentiation and myeloid-granulocytic hematopoiesis. Protocols, which aim at activating miR-221/222 expression in HSC may be helpful to maintain or re-establish hematopoietic potency through longevity, bone marrow homing capacity, quiescence and myeloid-lymphoid multipotency over myeloid-biased hematopoiesis.

## MATERIALS AND METHODS

### Mice

All mice were bred and kept in the breeding and experimental animal facilities of the Deutsches Rheumaforschungszentrum in Berlin Marienfelde and Berlin Mitte under SPF conditions. For all studies 6-12 weeks old mice were used. MiR-221/222^flox/flox^ mice on C57B6J background in which miR-221 and 222 were flanked by loxP sites were generated in the Helmholtz Zentrum München, Deutsches Forschungszentrum für Gesundheit und Umwelt (MAH-2166). Heterozygous C57BL6.Cg-Commd10^Tg(Vav1-icre)A2Kio/^J (The Jackson Laboratory) male mice were bred with miR-221/222^flox/flox^ females to generate miR-221/222^fl/y-Tg(Vav1-icre)^ miR-221/222 knock out males (deficient, Def.) in the F1 generation. Heterozygous C57BL6.Cg-Commd10^Tg(Vav1-icre)A2Kio/^J male mice were used as controls (proficient, Prof.). The Ly5.1 (CD45.1) male mice were obtained from Charles River and used in serial transplantation experiments. Mice were kept for at least 7 days in the experimental animal facility before they were taken in experiments or analysis. All of the experimental procedures complied with the “National Regulations for the Care and Use of Laboratory Animals”, approved by the Landesamt für Gesundheit und Soziales, Berlin (T0334/13, G0050-17).

### Social stress induction of mice

For short-term social stress-induced perturbation of the hematopoietic system the mice were used within 18-20 hours after the transport from the breeding facility in Berlin-Marienfelde to the experimental animal facility Berlin-Mitte. Separation of the experimental animals in the breeding facility, transportation and new housing in the experimental facility is a standardized protocol, and the separation, transportation and adaptation to the new environment are certified stress factors for experimental mice.

### Standardized methods used in the study

Bone marrow and spleen preparation, flow cytometry and cell sorting(78), serial transplantations, preparation of total RNA followed by quantitative PCR (qPCR) analysis are used as standard laboratory protocols (see Supplementary Methods). In order to inhibit continued “ex vivo/in vitro” stimulation of transcription during cell preparation, handling of the samples, which would influence the transcriptome analyses reported here (qPCR and single-cell RNA-sequencing (scRNA-seq)), femurs and tibia from sacrificed mice were collected and from then on kept in media containing 2µg/ml Actinomycin-D(25, 27). Six to eight week-old mice were used.

### Quantitation of miR-221/222 expression on single cell level

To measure the expression level of miR-221 and miR-222 in single HSC, MPP1 and MPP2 populations, we combined the protocols of Chen et al(79), Tang et al(80) and Petriv et al (38) with modifications (see Supplementary Methods).

### Single cell RNA-library preparation, sequencing and transcriptome profiling

The methods for single cell RNA-library preparation, sequencing and transcriptome profiling are standardized protocols described earlier(53, 81). Modifications related to the current study are described in Supplementary Methods.

### Data Sharing Statement

For original data, contact.

The single cell RNA-seq datasets are available through NCBI under BioProject ID XXX and GEO accession number GSEXXX, respectively (in progress). Source data are provided with this paper. Normalized scRNA-seq data are in **Supplementary Table** available with the online version of this article. The softwares used in this study are open source and listed in Supplementary Methods(48, 82–85).

## ACKNOWLEDGEMENTS

We thank Andreas Radbruch, DRFZ Berlin, and Nikolaus Dietlein and Hans-Reimer Rodewald, DKFZ Heidelberg, for critical reading of our manuscript. We thank the members of the animal breeding and experimental facility (Deutsches Rheuma-Forschungszentrum (DRFZ)) for their technical support and advice; the staff at the Max Planck Institute for Infection Biology Flow Cytometry Core Facility for expertise and instrument support; V.D. Dang, L. Bauer, and K. Lehmann for technical support and advice. P.K.J thanks the Alexander von Humboldt Foundation for a Postdoctoral Research Fellowship in 2017-2018. We thank the DRFZ’s core services for excellent support in technology and expertise. This work was supported by the Leibniz Collaborative Excellence Grant CHROQ-K121/2018.

## SUPPORTING INFORMATION

### METHODS

#### Bone marrow and spleen preparation

Bone marrow and spleen derived single cell suspensions were prepared as described earlier(78). In order to minimize a potential continued “ex vivo” stimulation of gene expression, e.g. of immediate early gene expression, which might occur during bone marrow cell preparation, cell handling of the samples, which might influence the transcriptome analyses reported here (qPCR and single-cell RNAseq) femurs and tibia from sacrificed mice were collected into tissue culture medium containing 2µg/ml Actinomycin-D. Actinomycin-D is an efficient RNA polymerase inhibitor and is used to minimize the uncontrolled “ex vivo” activation of gene expression changes as also found in hematopoietic cells(25, 27).

#### Flow Cytometry and cell sorting

We adhered to the guidelines for flow cytometry and cell sorting of hematopoietic stem and progenitor population and major lineage populations in the bone marrow and the spleen as described earlier(78). We used the nomenclature and gating strategy of BM derived stem and progenitor populations described by(5–7). All gating strategy, surface marker expression and antibodies used in the studies are listed in **Supplementary Fig 5A-G**.

#### Serial transplantation

For the first and second transplantations 100 CD45.2^+^ HSCs (miR-221/222-proficient or - deficient) were sorted and transplanted together with 1×10^6^ Ly5.1 carrier bone marrow into lethally irradiated (9,5Gy) CD45.1^+^ hosts (Ly5.1). Since ActD is toxic, it could not be used in these transplantation experiments to freeze the transcriptional state prior to transplantation. Hence, the scRNAseq results obtained with transplanted HSC and progenitors had to be done with “in vitro” stressed cells.

Mice were treated with 100mg/l Baytril 1 week before and for 2 weeks after transplantation. To follow the transplantation efficiency, 10µl tail vain blood was collected in heparinized tubes and the major lineage fractions were analyzed every 4 weeks until 16 weeks after transplantation by flow cytometry.

#### Preparation of total RNA

Total RNA was isolated from equal number of *ex vivo* sorted miR-221/222-proficient or deficient cells using TRIzol Reagent (Invitrogen) according to the manufacturer’s user guide. Isolated RNA was then quantified and qualified by Fragment Analyzer with the HS NGS Fragment Kit (1–6000 bp) (Agilent).

#### Real Time PCR analysis

RT-PCR was performed using SuperScript IV Reverse Transcriptase (Invitrogen) cDNA synthesis reaction and Oligo d(T)18 primer (Thermo Scientific) according to the user guide. Quantitive PCR was performed using QuantiTect SYBR Green PCR Master Mix (Qiagen) according to the user guide using following primer sets:

Fos F: 5’-GCCCAGTGAGGAATATCTGGA-3’, R: 5’-ATCGCAGATGAAGCTCTGGT-3’, and Hprt F: 5’- AAGCTTGCTGGTGAAAAGGA-3’, R: 5’- TTGCGCTCATCTTAGGCTTT-3’ as endogenous control.

Micro RNA expression on bulk sorted cells was measured by TaqMan real time PCR. All reagents were obtained from ThermoFischer Scientific. Measurements were analyzed using the Δ/ΔCT method relative to normalized miR-221 expression in pre-BI cells. Sno202 was used as a housekeeping gene. All qPCR reactions were performed in triplicates and originates from minimum 3 biological parallels.

#### Quantitation of miR-221/222 expression on single cell level

To measure the expression level of miR-221 and miR-222 in single HSC, MPP1 and MPP2 populations, we combined the protocols of(79),(80) and(38) with slight modifications. Briefly, the single cells were sorted in lysis buffer suitable for the RT reaction, then 3-plex (miR-221-, miR-222- and Sno202 specific) RT was carried out. Before the conventional simplex TaqMan qPCR, a 3-plexed cDNA pre-amplification PCR was done. We included external calibration standards consisting of 10-fold serial dilutions of synthetic miRNA oligonucleotides of 10^5^-10^0^ copies (**Supplementary Fig 6A,B**). Analysis of these standards revealed that assay efficiency varied considerably and that sensitivity ranged between 10^0^ and 10^2^ molecules per reaction based on no template control (NTC) (**Supplementary Fig 6C,D**). Pre-BI cells of Pax5^-/-^ mice expressing high levels of miR-221 and -222 were used as positive control while setting up the protocol. Variations in assay efficiency were found independent of multiplexing level and were observed even for serial dilutions of miRNA standards in single-plex reactions, therefore the Sno202 level of single cell samples not reaching the NTC were considered as sorting failure and excluded from the analysis (<10% of all sorted cells).

#### Single cell RNA-library preparation and sequencing

Single cell BM suspensions of unperturbed miR-221/222-proficient and deficient mice were stained for preparative cell sorting and MPP, CLP, lin^-^c-kit^+^Sca1^-^ and lin^-^c-kit^-^Sca1^-^ (5200 each) cells (**Supplementary Fig 5**) were applied to the 10x Genomics workflow for cell capturing and generation of scRNA gene expression (GEX) library using the Chromium Single Cell 3′ Library & Gel Bead Kit.

Due to the low cell counts of stem and progenitor cells, the bone marrow derived lineage positive (B220^+^, CD3^+^, CD4^+^, CD8^+^, CD11b^+^, CD11c^+^,CD19^+^, Gr1^+^, NK1.1^+^, TER119^+^) cells were first depleted on magnetic column (LS from Miltenyi) and the remaining lin^-^ single cell suspensions from 4-4 perturbed or unperturbed miR-221/222-proficient and deficient mice were labeled separately for preparative FACS (**Supplementary Fig 5**), followed by staining with TotalSeq™-C anti-mouse Hashtag-1 or-4 (miR-221/222-proficient, perturbed: #1 or unperturbed: #4) or -Hashtag-2 or-5 (miR-221/222-deficient perturbed: #2 or unperturbed: #5) antibodies (BioLegned, #1: 155861, #2: 155863, #4: 155867, #5 155869). The unperturbed or perturbed miR-221/222-proficient samples were pooled with deficient samples for scRNAseq. 11584 HSC, 12605 MPP1 and 18721 MPP2 sorted from pooled unperturbed samples or 8900 HSC, 12400 MPP1 and 9100 MPP2 sorted from pooled perturbed samples were then applied to 10x Genomics workflow for cell capturing and scRNA gene expression (GEX) library preparation using the Chromium Single Cell 5′ Library & Gel Bead Kit as well as the Single Cell 5′ Feature Barcode Library Kit (10x Genomics). After cDNA amplification the CiteSeq libraries were prepared separately using the Single Index Kit N Set A while final GEX libraries were obtained after fragmentation, adapter ligation, and final Index PCR using the Single Index Kit T Set A.

The bone marrow derived lineage positive cells were first depleted on magnetic column and the remaining single cell suspensions from four, 2-times transplanted animals were stained for preparative sorting of CD45.2^+^ cells. After sorting 7045 HSC, 6363 MPP1 and 3532 MPP2 from miR-221/222-proficent HSC transplanted mice, the cells were separately applied, while 1780 HSC, 771 MPP1 and 422 MPP2 sorted from deficient HSC transplanted mice were pooled after sorting and applied to the 10x Genomics workflow using the Chromium Single Cell 5′ Library & Gel Bead Kit and Single Index Kit T Set A.

For all libraries prepared the fragment sizes were determined using the Fragment Analyzer with the HS NGS Fragment Kit (1–6000 bp) (Agilent) and library concentrations were determined with Qubit HS DNA assay kit (Life Technologies).

3′ GEX libraries and 5′ GEX+CITE libraries were sequenced on a NextSeq500 device (Illumina) using High Output v2 Kits (150 cycles) or on a NextSeq2000 device (Illumina) using either P2 reagents (200 cycles) or P3 reagents (200 cycles or 100 cycles) with the recommended sequencing conditions for (read1: 26nt, read2: 98nt, index1: 8nt, index2: n.a.).

#### Single-cell transcriptome profiling

Raw signals were demultiplexed and converted to fastq files using DRAGEN (Ilumina). Detection of intact cells and expression quantification was performed by cellranger (version 5.0.0) using count in default parameter settings with number of expected cells set to 3000 and refdata-cellranger-mm10 as reference. Further analysis was done in R (version 4.1.2) using the Seurat package (version 4.0.5).

The integration of the datasets from HSC, MPP1, MPP2, MPP(3–4), CLP, lin^-^c-kit^+^Sca1^-^ and lin^-^c-kit^-^Sca1^-^ cells of unperturbed miR-221/222-proficient and deficient mice was performed by following the integration pipeline as described in the FindIntegrationAnchors (Seurat) R Documentation. Firstly, each library was log-normalized using NormalizeData, 2000 variable genes were detected using vst as selection method with FindVariableFeatures, variable genes were scaled using ScaleData and 50 principle components were computed using RunPCA. Next, common anchors were identified by FindIntegrationAnchors using rpca as reduction, 2000 anchor features and 1:30 dimensions. Finally, libraries were merged using IntegrateData. The integrated data was further analyzed, by a uniform manifold approximation and projection (UMAP) sequentially using ScaleData, RunPCA for 50 principle components and RunUMAP using 1:30 principle component. Transcriptionally similar clusters (T-clusters) were identified by shared nearest neighbor (SNN) modularity optimization with FindNeighbors using 1:30 principle components and FindClusters with the resolutions of 0.2. The UMAP was evaluated by projection of scores for selected developmental stages, defined as the sum of log-normalized expression values.

In the aggregated dataset we detected a total 15 distinct clusters of BM lin^-^ populations, (called total-(T-) clusters). Excluding clusters representing less then 5% of all analyzed cells, we further investigated T-cluster 0 to 11. T-clusters 0, 2 and 10 could be characterized by the expression of HSC-related genes, mostly from sorted HSC. T-clusters 3 and 4 had accumulated expression of cell cycle-active genes – cluster 3 for G1/S-phase and cluster 4 for G2/M phase, mostly from sorted MPP1-4 cells. T-cluster 5 contained T and NK lymphoid directed, T-cluster 6 contained B cell directed cells, mostly from sorted CLP. T-cluster 8 contained erythropoiesis-directed, T-cluster 9 megakaryocyte-platelet-directed genes and T-cluster 1 had accumulated genes of earlier phases of myelopoiesis, mostly from sorted lin^-^c-kit^+^Sca^-^ or lin^-^c-kit^-^Sca^-^ cells. Finally, T-cluster 7 expressed genes of more mature stages of basophil and mast cell development, while in T-cluster 11 genes expressed in more mature stages of neutrophils and granulocytes were predominant. These T-clusters 7 and 11 consisted mostly sorted lin^-^ c-kit^+^Sca^-^ or lin^-^c-kit^-^Sca^-^ cells.

The same workflow was used to merge HSC, MPP1 and MPP2 datasets. In particular, the HSC, MPP1 and MPP2 from unperturbed and perturbed miR-221/222-proficient and deficient mice, HSC, MPP1 and MPP2 sorted after the second transplantation of miR-221/222-proficient HSC as well as the pool of HSC, MPP1 and MPP2 sorted after the second transplantation of miR-221/222-deficient HSC were integrated, a UMAP was computed and transcriptional cluster were defined using the resolution of 0.6. Clusters (E-clusters) with low quality were identify by visual inspection of UMI counts, number of expressed and percentage of mitochondrial genes. The contaminating cluster with erythrocytes contained cells with high Hbb expression and higher UMI counts and number of detected genes. Cells from low quality clusters as well as the contaminating cluster with less then 5% contribution to all cells were removed from further analysis. Libraries with hashtag-labeled antibody stainings of miR-221/222-proficient and deficient cells were demultiplex by contrasting arcsinh transformed hashtag 1 or 4 and hashtag 2 or 5 in a scatterplot and manual gating.

Differential gene expression analysis and comparison of number of expressed genes and UMIs per cell, were performed based on down sampled libraries. In analogy to DESeq2 library size normalization, proposed for bulk sequencing(86), pseudo bulk samples were created for each library after removal of low quality and contaminating clusters by summing up the read counts for each gene. Next, the geometric mean was calculated for each gene expressed in at least 75% of the considered cells in each library and at least 75% of miR-221/222-proficient and deficient cells in the pools. The library size factors were defined as the median ratio between the UMI counts of the library and the corresponding geometric mean normalized to the number of considered cells. Down sampling was performed by subsampling of UMIs with sampling rates defined as the ratios of the minimal and the respective size factor. Numbers of detected genes and UMIs were computed after resembling the gene counts. Gene expression represents log2p transformed UMI counts. Differential expression analysis was performed based on subsampled but not normalized values. In particular, for each gene log2p transformed, all 0 values removed and a Mann-Whitney-Test was performed. Next, genes were ranked by the fraction of expressing proficient (for miR-221/222 sensitive genes) or unperturbed (for perturbation sensitive genes) cells and a linear regression was performed on the log2p fold change between miR-221/222-deficient and proficient or fold change between perturbed and unperturbed samples (mean log2Exp_deficient_-mean log2Exp_proficient_ or mean log2Exp_perturbed_-mean log2Exp_unperturbed_). The linear regression was performed using Prism (Version 9.0). Genes were defined as differentially expressed, if found in at least 30% of the cells and the log2 fold change exceeding the 99% prediction band (PB).

#### Code availability

The softwares used in this study are open source. Cellranger from 10xgenomics: (https://support.10xgenomics.com/single-cell-gene-expression/software/downloads/latest) Seurat packages 4.1.1: (https://cloud.r-project.org/web/packages/Seurat/index.html). Used scripts are available upon request. (https://CRAN.R-project.org/CRANlogo.png; https://cloud.r-project.org/web/packages/Seurat/index.html) CRAN-Package Seurat: (https://cloud.r-project.org/web/packages/Seurat/index.html), (cloud.r-project.org). A toolkit for quality control, analysis, and exploration of single cell RNA sequencing data. ‘Seurat’ aims to enable users to identify and interpret sources of heterogeneity from single cell transcriptomic measurements, and to integrate diverse types of single cell data. See (82–85) for more details.

## SUPPLEMENTARY/ONLINE TABLE

**Supplementary Table 1.** Normaliezed scRNA-seq data used in the study.

## SUPPLEMENTARY FIG CAPTIONS

**Supplementary Fig 1: Flow cytometry analysis of unperturbed and in vivo social stress-perturbed miR-221/222-proficient and deficient hematopoietic cells in bone marrow and spleen:** (**A,B**) Analysis of flow-cytometry measurements on unperturbed and social-stress perturbed bone marrow and spleen derived cells. (**A**) Single cell suspensions of 2 tibia and femurs of mice or (**B**) from spleens of unperturbed miR-221/222-proficient (open squares) and miR-221/222-deficient (closed squares) mice or perturbed miR-221/222-proficient (open triangles) or miR-221/222-deficient (closed triangle) mice were prepared in matched pairs, analyzed with flow-cytometry and the numbers of cells were plotted. Red lines indicate the mean values. One-way ANOVA with Tukey post-test was used to evaluate statistical significance, exact *p* values are given above the comparing lines.

**Supplementary Fig 2: Differentially expressed genes upon in vivo social stress-perturbation in MPP1 and MPP2 cells.** Genes with higher expression after short-term perturbation (red dots and gene symbols are selected genes) in (**A**) MPP1 and (**B**) MPP2 cells are presented by comparative differential expression analysis of unperturbed versus perturbed cells. Differentially expressed genes are above the significance limit (red dashed). The log2 fold-change expression values were plotted against the numbers of unperturbed cells expressing the gene. The right side of the green line indicates genes expressed in more than 30% of the cells.

**Supplementary Fig 3: G1/S and G2/M cell cycle phases on UMAPs of early (E) and total (T) hematopoietic compartments. (A)** The integrated data of miR-221/222-proficient and deficient unperturbed, short-term perturbed and serial transplanted HSC, MPP1 and MPP2 populations after single cell transcriptome sequencing (Early) or (**B**) HSC, MPP1, MPP2, MPP, CLP, lin-ckit+Sca1- and lin-ckit-Sca1-populations (Total) were further analyzed, by a uniform manifold approximation and projection (UMAP). Cells with characteristic gene expression pattern for G1/S (left) or G2/M (right) cell cycle-related genes are shown on a composite picture for proficient and for deficient cells. For plotting, gene-set modules of G1/S and G2/M genes were used (**Supplementary Table 1**).

**Supplementary Fig 4.: MiR-221/222 expression in hematopoietic cells.** (**A**) Relative miR-221 expression in bulk sorted hematopoietic progenitor, B-cell subsets, NK cells, granulocytes and T cells in bone marrow and thymus (n=5 mice). Expression of miR-221 in the different bulk sorted populations are displayed as fold change relative to miR-221 expression in preBI cells. (**B**) Relative miR-221 expression in bone marrow of miRNA proficient and deficient LSK and MPP subsets (n=5 mice). HSC/MPP1: pool of hematopoietic stem cell and multipotent progenitor (MPP)1, MPP3-4: pool of MPP3 and MPP4 populations, CLP: common lymphoid progenitor, PreBI: precursor BI cell, Pro-Pre-B: progenitor of precursor B cell, NK cell: natural-killer cells. LSK: bone marrow derived Lin^-^Kit^+^Sca1^+^ population.

**Supplementary Fig 5: Gating strategy and surface marker expression of stem and progenitor populations in the bone marrow and lineage cells in the spleen.** After preparing single cell suspensions of (**A**) unperturbed or (**B**) perturbed miR-221/222-proficient, or of (**C**) unperturbed or (**D**) perturbed miR-221/222-deficient bone marrow, stem and progenitor cells were stained for flow-cytometry analysis. (**E**) Single cell suspension of miR-221/222-proficient or deficient spleen or blood was prepared and the major hematopoietic lineage cells were stained for flow-cytometry analysis. (**F**) Surface marker expression of the measured hematopoietic stem, progenitor and lineage cells. HSC were gated on lin^-^ (B220^-^CD3^-^CD4^-^CD8^-^CD19^-^CD11c^-^CD11b^-^Gr1^-^NK1.1^-^ TER119^-^) c-kit^+^Sca1^+^Flk2^-^CD34^-^CD150^+^CD48^-^ cells. MPP1 was gated on lin^-^c-kit^+^Sca1^+^Flk2^-^ CD34^+^CD150^+^CD48^-^ cells. MPP2 were gated on lin^-^c-kit^+^Sca1^+^Flk2^-^CD34^+^CD150^+^CD48^+^ cells. MPP3 were gated on lin^-^c-kit^+^Sca1^+^Flk2^-^CD34^+^CD150^-^CD48^+^ cells. MPP4 were gated on lin^-^c-kit^+^Sca1^+^Flk2^+^CD34^+^CD150^-^ CD48^+^ cells. CLP were gated on lin^-^c-kit^lo^Sca1^lo^Flk2^+^IL7R^+^ cells. CMP were gated on lin^-^c-kit^+^Sca1^-^ CD34^+^CD16/32^-^ cells. MEP were gated on lin^-^c-kit^+^Sca1^-^CD34^-^CD16/32^-^ cells. GMP were gated on lin^-^c-kit^+^Sca1^-^ CD34^+^CD16/32^+^ cells. CD4 T cell were gated as CD3^+^CD4^+^, CD8 T cell were gated as CD3^+^CD8^+^ cell. B cells were gated as CD4^-^CD8^-^B220^+^CD19^+^ cell. Myeloid cells were gated as CD4^-^CD8^-^B220^-^CD19^-^CD11b^+^Gr1^-^ cells. Granulocyte (Gran.) were gated on CD4^-^CD8^-^B220^-^CD19^-^CD11b^+^Gr1^+^ cells. (**G**) Antibodies used in the analyses.

**Supplementary Fig 6: Strategy for the detection of copy numbers of miR-221/222 molecules in single cells.** (**A**) 10e5-10e0 copies of serially diluted synthetic miR-221 oligonucleotide was measured 3 times (Run1-3) in 3 technical replicates. The TaqMan qPCR measurements were done after or without (without pre-amp) 10 cycle PRC amplification of the reverse transcript. (**B**) 10e5-10e0 copies of serially diluted synthetic miR-222 oligonucleotides were measured 2 times (Run1 and 2) in 3 technical replicates. The TaqMan qPCR measurements were done after 10 cycle PRC amplification of the reverse transcript. (**C**) 45 single-cell sorted miR-221/222- proficient Lin^-^c-kit^+^Sca1^+^CD150^+^CD48^-^ (pool of HSC and MPP1) cells were directly sorted in lysis buffer, suitable for miR-221 or miR-222-specific reverse transcription reaction. After 10 cycle PRC amplification of the reverse transcript, the copy numbers of miR-221 (red) and miR-222 (blue) were determined in individual cells using standard curves developed in the same reaction plate. Red and blue dots indicate technical replicates (**D**) The limit of detection in miR-221 and -222 copy numbers were assigned by the highest value of 10 no-template controls (NTC), where no cell was sorted in the lysis buffer. Red and blue dots indicate technical replicates. Pax5^-/-^ pre-BI cells were used as positive control. The mean of the technical replicates are calculated and plotted for 10 individual cells.

